# Variation in insect herbivore communities on individual plants reveals phylogenetic signal in uncertainty of attack in Brassicaceae

**DOI:** 10.1101/2020.12.06.413724

**Authors:** Daan Mertens, Klaas Bouwmeester, Erik H. Poelman

## Abstract

As a result of co-evolution between plants and herbivores, related plants often interact with similar communities of herbivores. On individual plants, typically only a subset of interactions is realized. The stochasticity of realized interactions leads to uncertainty of attack on individual plants and is likely to determine adaptiveness of plant defence strategies. Here, we show that across 12 plant species in two phylogenetic lineages of the Brassicaceae, variation in realized herbivore communities reveals a phylogenetic signal in the uncertainty of attack on individual plants. Individual plants of Brassicaceae Lineage II were attacked by a larger number of herbivore species from a larger species pool, resulting in a higher uncertainty of realized antagonistic interactions compared to plants in Lineage I. We argue that uncertainty of attack in terms of realized interactions on individual plants is ecologically relevant and must therefore be considered in the evolution of plant defences.

## Introduction

Over their lifetime, plants interact with numerous organisms of which many are herbivorous. In addition to a mixture of environmental effects such as host plant community composition and abiotic conditions, the set of antagonistic interaction partners of a plant species is strongly determined by selection pressures and phylogenetic history (Morales-Castilla et al. 2015, Hutchinson et al. 2017). Insect herbivores may exert selection that promotes novel defence strategies in plants and these events often coincide with radiation of herbivore species that have counteradaptations to these novel defences (Edger et al. 2015). Even though the relationship between plant phylogenetic distance and overlap in herbivore communities is variable across plant clades, co-evolutionary processes often result in non-random structuring of plant-associated antagonist communities (Bergamini et al. 2017, Rapo et al. 2019, Cirtwill et al. 2020).

However, these macro-evolutionary patterns are generally described for the full potential pool of antagonist species interacting with a plant species, whereas natural selection by these antagonists acts on the level of plant individuals. In many plant species, there is a discrepancy between the full potential of antagonistic interaction partners at the plant species level and the subset of realized interactions for individual plants (Lewinsohn et al. 2005, Kuppler et al. 2016). The subset of antagonists actually colonizing individual plants is determined by intraspecific variation in plant traits (Barbour et al. 2015, Barbour et al. 2018), variation in functional traits across antagonist species or individuals (Zytynska and Preziosi 2013), priority effects in community assembly in which a first colonizer affects the likelihood of attack by other antagonists (Lill and Marquis 2003, Stam et al. 2018), environmental heterogeneity and habitat filtering (Johnson and Agrawal 2005, Agrawal and Fishbein 2006, Violle et al. 2012), as well as stochastic processes (Shoemaker et al. 2020). Moreover, plant species may differ in the relative proportion of realized interactions per individual plant out of the full antagonist species pool and thus differ in the uncertainty of which antagonists attack individual plants. For example, plant species that interact with a larger species pool of herbivores face a larger number of potential herbivore species at the level of individual plants, possibly resulting in a larger uncertainty of realized interactions. At the same time, the uncertainty of interactions experienced by individual plants is determined by both the proportion as well as the absolute number of realized interactions and dominance of interactions by key herbivore species. The uncertainty of attack on individual plants is a major component of theories on plant defence plasticity (Poelman and Kessler 2016, Salazar et al. 2016b, Karban 2019). Plants evolved inducible defences that are activated upon recognition of attack to save metabolic costs associated with production and maintenance of defence when herbivores are absent (Meldau et al. 2012). Plasticity in defence also allows plants to tailor defences to specific antagonists and to integrate responses to multiple attackers (Van der Ent et al. 2018). Variation across plant species in the uncertainty of attack may strongly impact the evolution of such plastic defence strategies. It is to be expected that related plant species face more similar levels of uncertainty in their interactions as closely related plant species tend to interact with herbivore communities that are similar in composition (Agosta 2006, Agrawal and Fishbein 2006, Futuyma and Agrawal 2009). However, it is unknown whether phylogenetic signals in uncertainty of community composition or structure on individual plants exist. Understanding how the overall uncertainty of interactions that plants experience varies within and among plant species can provide valuable insights in the evolutionary history of defence strategies (Violle et al. 2012, Ohgushi 2016, Salazar et al. 2016a, Stam et al. 2018).

To determine phylogenetic signals in plant-herbivore community composition and explore variation in uncertainty of realized interactions across plant individuals, we compare insect herbivore community characteristics among 12 Brassicaceae species, belonging to two major phylogenetic lineages (Lineage I and Lineage II) within this family. We hypothesize that the composition of the herbivore community with which plants interact correlates with plant phylogeny, and that this phylogenetic signal is retained in the uncertainty of realized interactions on individual plant level. We explicitly test whether plant phylogenetic distance predicts i) the species richness, diversity and the proportion of realized interactions on individual plants out of the full potential herbivore community as a measure of uncertainty of attack on individual plants, and ii) the similarity in the average herbivore community composition and structure on individual plants within a species. We subsequently test whether the phylogenetic signal correlates with similarity in plant growth traits. We then evaluate which evolutionary models best predict phylogenetic signals in herbivore community diversity and uncertainty of attack in Brassicaceae and discuss its consequences for the evolution of plant defences.

## Results

### Plant phylogeny structures the number and proportion of realized interactions on individual plants

By repeatedly monitoring insect herbivore communities during the life span of individual plants of 12 annual Brassicaceae species (Supplementary File 1 Table A, Supplementary Fig. 1), we identified that the network of insect herbivores interacting with the 12 plant species was strongly connected (Fig. 1). Overall, both the interaction network based on herbivore incidence, as well as its abundance-based equivalent were characterized by high levels of connectance and nestedness and low levels of specialization (Table 1, Supplementary File 2 Tables A – D). The two phylogenetic lineages showed no distinct subgroups of interactions and the full insect herbivore species pools were largely shared among all plant species (Fig. 1). Plant species in our network showed significantly higher levels of diversity in their interactions with herbivore species compared to expectations inferred by two separate null models. However, overall level of plant specialization remained low, and was dependent on the subset of the herbivore community under investigation. Plant-species-specific levels of specialization in their interactions with herbivores (expressed by Blüthgens’ di’) were highest for the abundance-based network analysis of the sap-feeding herbivores (Supplementary File 2 Table D). The increased specialization of plant interactions with this community subset is likely driven by the exponential increase in population size of aphids on plants they successfully colonize, mimicking patterns of increased interaction specialization (Supplementary File 2 Table E). Although plant species differed in the diversity of their full herbivore communities, these patterns were generally not phylogenetically structured (Table 1, Supplementary File 3 Tables A – B). The most apparent plant phylogenetic signal in overlap of the species pool of insect herbivores was found for a higher species richness of sap-feeding herbivores on plant species of Lineage II compared to plant species of Lineage I (LM: df = 1; *F* = 10.7390; *P* = 0.0083). However, the full herbivore species pools associated with Brassicaceae species of Lineage II were not statistically different in size or diversity compared to herbivore communities associated with species of Lineage I. None of the abundance-based diversity measures (i.e. Shannon and Simpson indices) calculated for the community subset of sap-feeding herbivores, nor the diversity measures calculated for the subset of chewing herbivores were significantly different among lineages (Supplementary File 3 Table B).

**Fig. 1.**
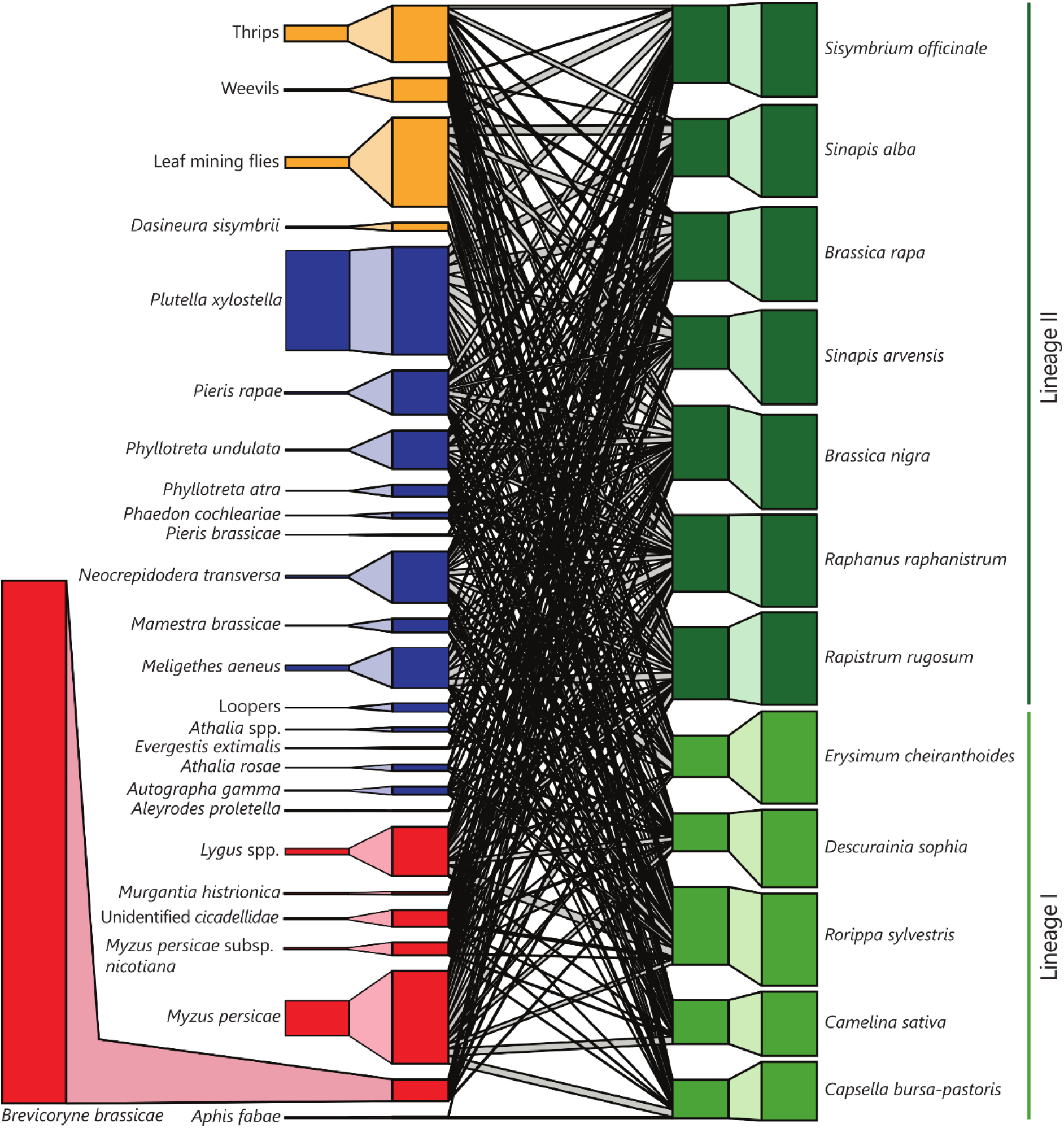
Interactions between Brassicaceae and their herbivore communities are characterized by low levels of specialization. Bipartite network of the total plant – herbivore community observed in our common garden experiment. Boxes on the outside of the diagram represent relative abundances of herbivores and plants. Boxes on the inside of the diagram represent interaction frequencies adjusted for uneven sampling of plant individuals (i.e. incidence-based network). Shaded areas between boxes on the outside of the diagram and on the inside of the diagram (within the same trophic level) depict the association between relative abundance and standardized prevalence in the field. Lines connecting herbivore species and plant species represent realized interactions, and width of these lines represent the relative number of (incidence) interactions. Colours of boxes represent sap-feeding herbivores (red), chewing herbivores (blue), and unclassified herbivores (yellow) on the trophic level of herbivores, and Brassicaceae species belonging to Lineage I (light green), and Lineage II (dark green) at the trophic level of plant species.

**Table 1.** Overview of the herbivore community matrices used in our analyses, presenting the biological level and interpretation of values in the matrix.

| Biological level | Values represent | Standardisation | Transformation | Used in analysis |
| --- | --- | --- | --- | --- |
| A. Plant individual | Cumulative abundance of herbivores | - | - | Univariate analysis of Shannon and Simpson indices |
| B. Plant individual | Incidence of herbivores | - | - | Univariate analysis of species richness, Uni- and Multivariate analysis $\beta$ diversity, Multivariate analysis of community composition |
| C. Plant individual | Cumulative abundance of herbivores | - | $\log(x + 1)$ | Multivariate analysis of community structure, Multivariate regression of plant traits, Phylogenetic similarity analysis. |
| D. Plant species | Cumulative abundance of herbivores | - | - | Network analysis |
| E. Plant species | Percentage of plants | Proportion | - | Network analysis |

The proportion standardisation (matrix E) was calculated by summing the number of plants belonging to a specific plant species on which the herbivore species occurred and dividing this number by the total number of plants belonging to this species in our experiment. We chose not to standardize the abundance-based associational matrix D as the relation between the summed abundance of herbivores and plant sample size is likely to be herbivore-species-specific due to differences in the biology of the herbivores

Importantly, in contrast to the absence of a strong plant phylogenetic signal in the full pool of insect herbivores associated with plant species, the realized interactions with herbivores on individual plants were clearly structured by plant phylogenetic lineage. Individual plants of species in Lineage II were attacked by a more species-rich herbivore community than plants of species belonging to Lineage I (Fig. 2, Table 1, Supplementary File 3 Tables C – D). The higher species richness was not dependent of the herbivores’ feeding guild, showing that individual plants in Lineage II interacted with a more species-rich chewer community as well as a more species-rich sap-feeding herbivore community (Supplementary Fig. 2). A marginally lower proportion of potential interactions was realized on individual plants of Lineage II than Lineage I (expressed by Whittaker’s β diversity; Anderson et al. 2011) (LMM: df = 1; *χ*2 = 4.4132; *P* = 0.0357) (Fig. 2). As the proportion of realised interactions was comparable across the two plant lineages and richness of realised as well as potential interactions was higher for plants in Lineage II, plants of Lineage II were characterised by a higher uncertainty in the realized interactions they encounter. Across all 12 brassicaceous species, uncertainty of attack as illustrated by variation in herbivore community composition on individual plants was generally high and primarily caused by differences in the identities of herbivores rather than variation in the number of herbivore species attacking individual plants (Table 1, Supplementary Fig. 3, Supplementary File 4 Table A). Plant species across lineages varied significantly in the average values of both the Shannon and Simpson indices, identifying differences in evenness of community structure across plant species (Supplementary Figs 4–5, Supplementary File 3 Table C). Except for a significantly higher Shannon index associated with chewing herbivore communities on plant species belonging to Lineage II, these indices were not significantly different between the two lineages (Supplementary File 3 Table D).

**Fig. 2.**
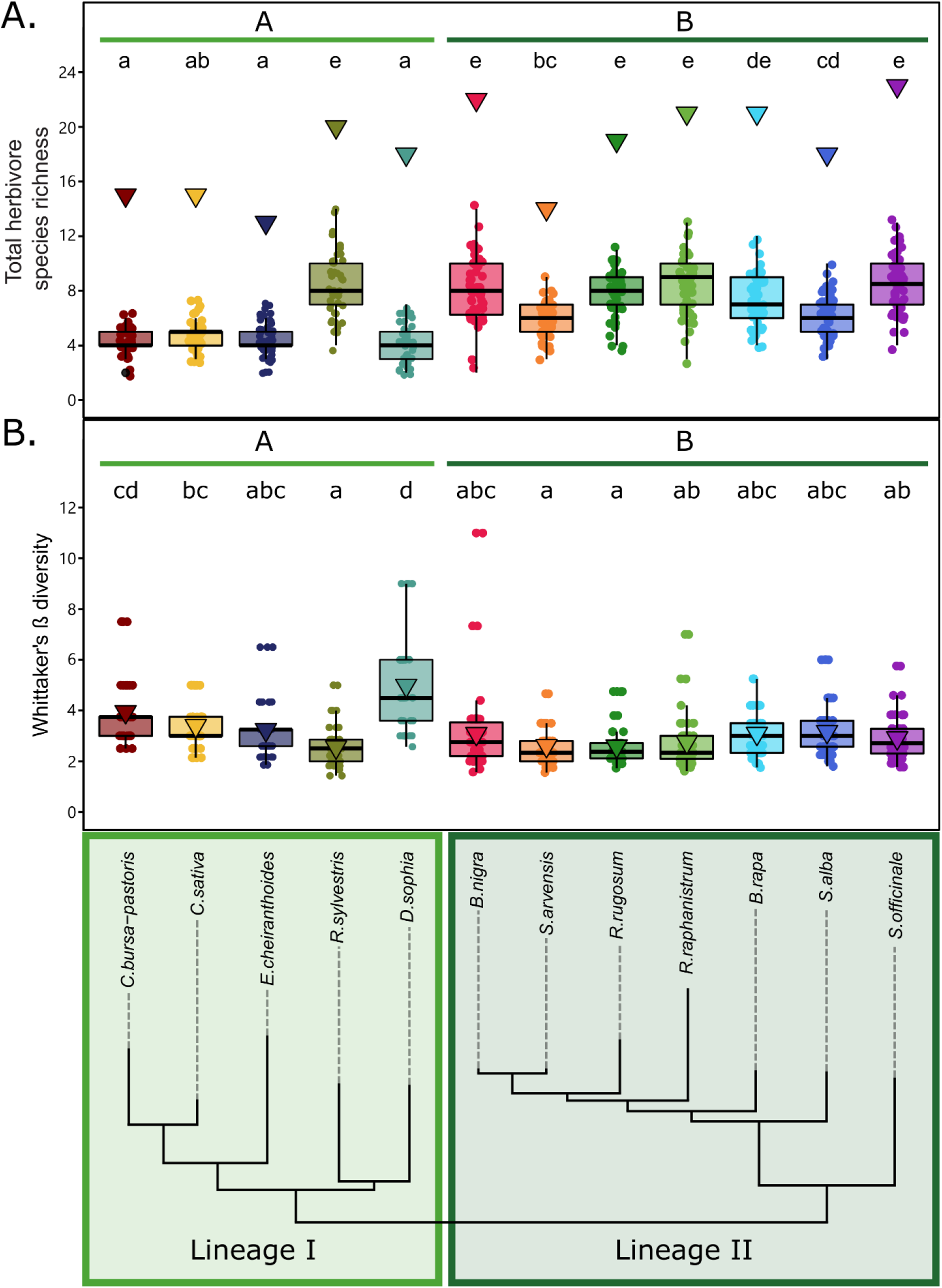
Herbivore diversity and uncertainty of attack on individual plants is dependent on plant species and structured by plant phylogeny. (A) Herbivore species richness on individual plants correlates with plant phylogenetic lineages and (B) the relation between the herbivore species richness and the number of realized interactions (Whittaker’s β) as measures for uncertainty of attack on individual plants observed across plant phylogenetic lineages (lower panel). Triangles depict the total number of herbivore species and the average β diversity associated with the total number of individuals per plant species respectively. Dots represent the number of herbivore species or the β diversity observed on individual plants of the respective plant species. Box-whiskers summarize the variation in observations at the level of plant individuals. Statistical analyses were performed by applying Linear Mixed Models (LMM) with species or phylogenetic lineage as explanatory factors and including plot and, when estimating the diversity for phylogenetic lineages, plant species as random factors in our models. To account for heterogeneity of variance, we allowed the variance to be different for the different species or lineages in our model. Different letters indicate significant different means (*P* < 0.05), adjusted for multiple testing by Tukey HSD. Significant differences across lineages (plant species grouped by the coloured horizontal bars) are indicated with capital letters. Statistical analyses were performed separately for the different panels.

Overall, these results indicate that individual plants of Lineage II were exposed to higher uncertainty of attack than plants of Lineage I, and that this uncertainty is mainly driven by the high absolute richness of potential species interactions and the high number of realized interactions per plant individual.

### Individual plants of different species differ in herbivore community composition and structure

The average composition of herbivore communities on individual plants differed between plant species and between phylogenetic Lineage I and II (Fig. 3 A – B, Table 1 – 2). These differences were emphasized when taking the community structure into account (Fig. 3 C – D, Tables 1 – 2). The plant phylogenetic signal in herbivore communities on individual plants was further supported by a significant correlation between the dissimilarity in community composition and structure among individual plants and their phylogenetic dissimilarities at the species level (Fig. 4, Table 3). Pairwise comparisons of realized communities on individual plants were almost always significantly different between plant species (Supplementary File 4 Table B – C). Overall, these results indicate that plant species and phylogenetic lineages were characterized by the average herbivore community associated with individual plants (Fig. 4). We show that plants belonging to different species differ in the community of herbivores they interact with and that this difference in herbivore communities is significantly larger for more distantly related plant species.

**Fig. 3.**
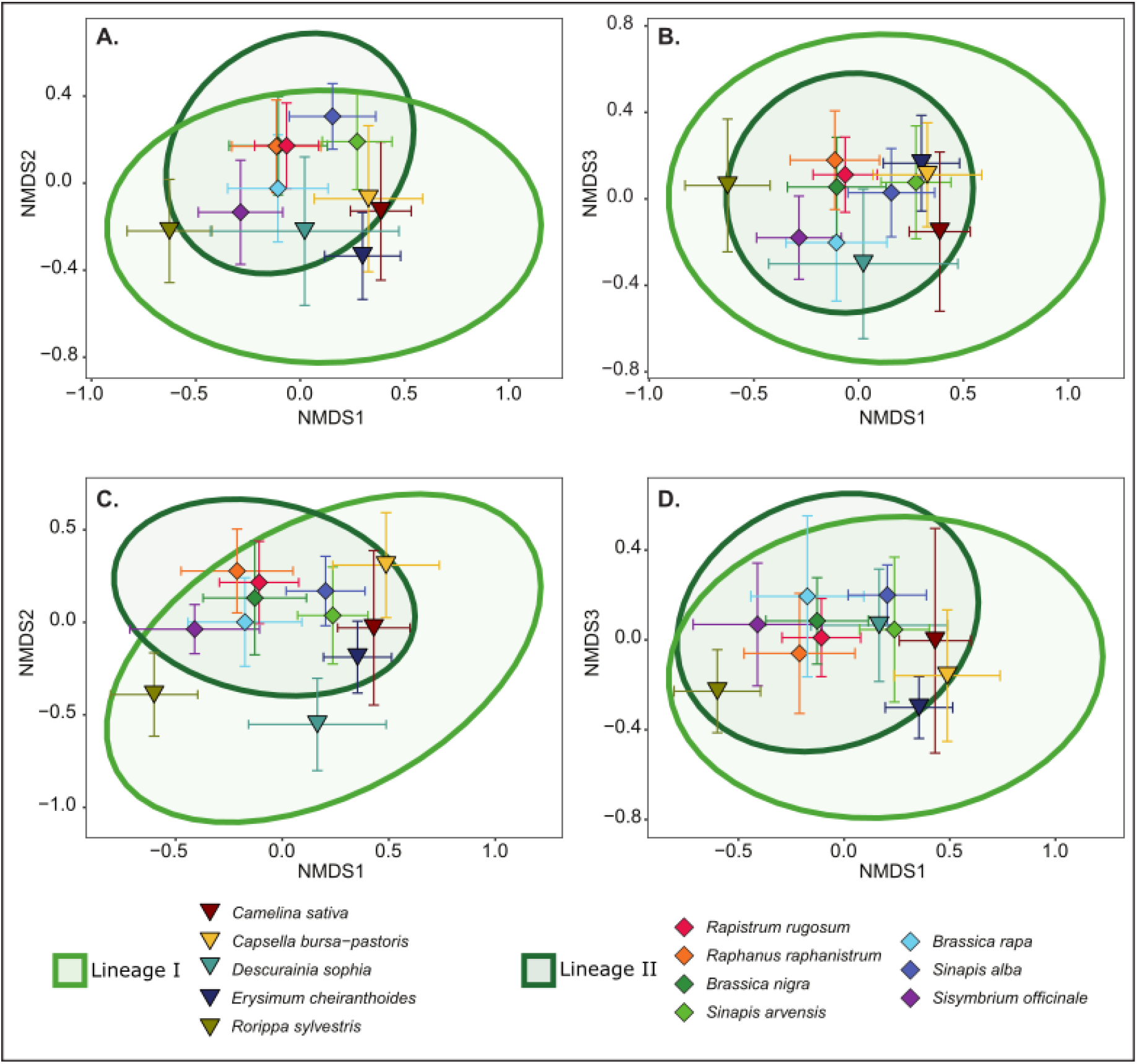
Composition and structure of herbivore communities differ across plant species but overlap across plant lineages. Ordination of observed herbivore community composition (expressed by incidence of herbivores, panels A and B), and structure (expressed by log (x + 1) transformed cumulative herbivore abundance data, panels C and D), according to three NMDS ordination axes (stress = 0.18 and 0.19 respectively). Triangles and diamonds represent the centroid of the variation in communities associated with plants belonging to Lineage I and Lineage II respectively and are coloured according to plant species. Error bars around the plant centroids represent the 95% confidence interval around the estimation of the mean. Ellipses are coloured according to plant lineage and depict the 95% interval of a multivariate t-distribution around the centroids of each of the two plant lineages.

**Table 2.** Differences between plant lineages or plant species in the composition or structure of the herbivore community associated with plants estimated by PERMANOVA analysis.

| Community subset | Biological level | Composition |  |  | Structure |  |  |
| --- | --- | --- | --- | --- | --- | --- | --- |
|  |  | Pseudo- |  |  | Pseudo- |  |  |
|  |  | df | *F* | *P* | df | *F* | *P* |
| Full herbivore community | Lineage | 1 | 3.2124 | **0.0070** | 1 | 12.735 | **0.0051** |
|  | Plant species | 11 | 23.936 | **0.0001** | 11 | 23.857 | **0.0001** |
| Chewing herbivore community | Lineage | 1 | 4.9949 | **0.0050** | 1 | 9.8834 | **0.0104** |
|  | Plant species | 11 | 24.15 | **0.0001** | 11 | 21.165 | **0.0001** |
| Sap-feeding herbivore community | Lineage | 1 | 0.7023 | 0.4925 | 1 | 7.6642 | **0.0198** |
|  | Plant species | 11 | 16.4718 | **0.0001** | 11 | 17.402 | **0.0001** |

Community composition was analysed by incidence observations, and community structure assessed by analysing log (x + 1) transformed cumulative abundance observations. The analysis was performed separately for the full herbivore community, the chewing herbivore community and the sap-feeding herbivore community associated with plants. Significant *P* values (*P* < 0.05) are indicated in bold and are assessed by 999 random permutations, while taking the dependency of observations in our experiment into account.

**Table 3.**
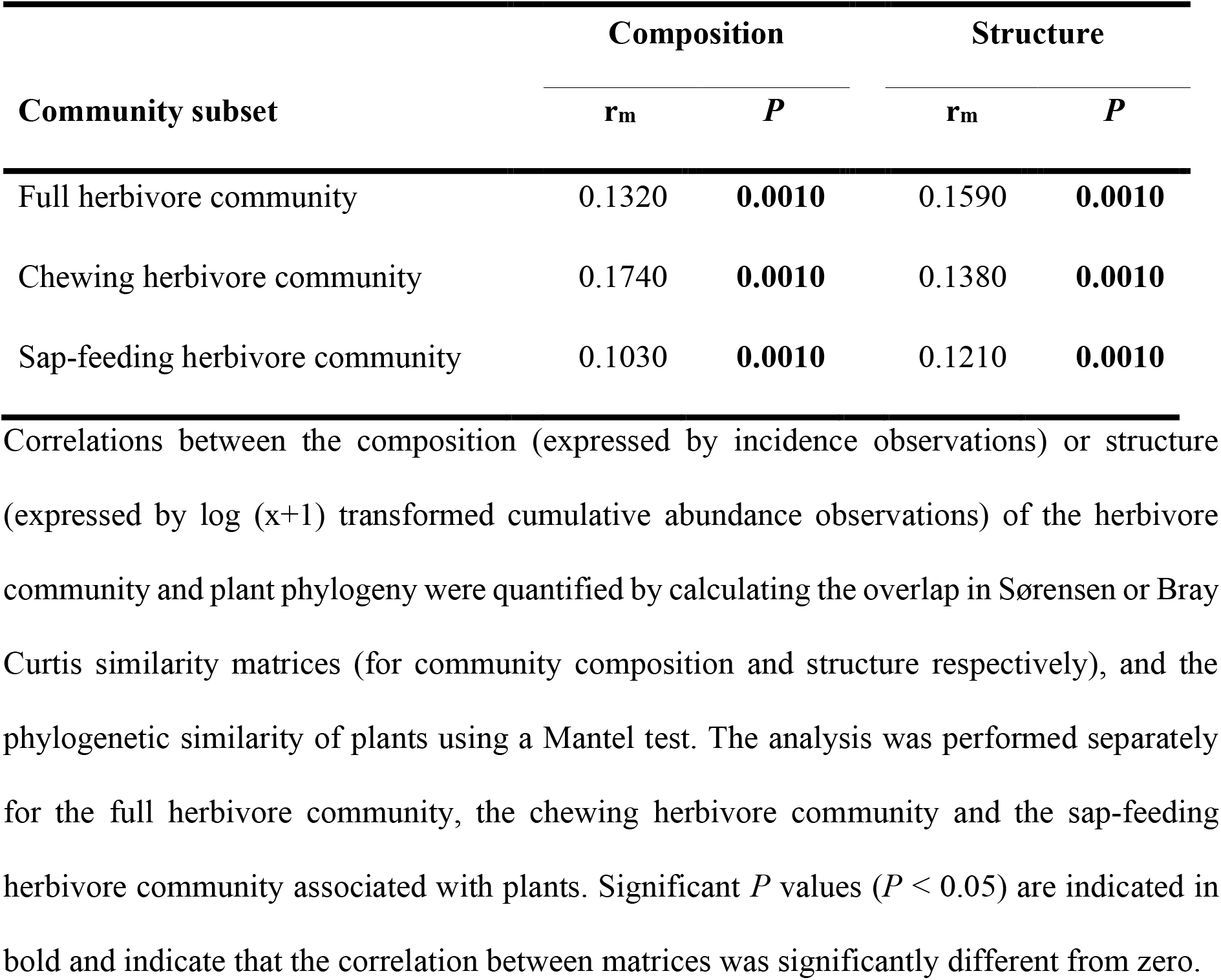
Correlation between the herbivore community composition or structure of plants and their phylogenetic similarity inferred from ITS sequences.

**Fig. 4.**
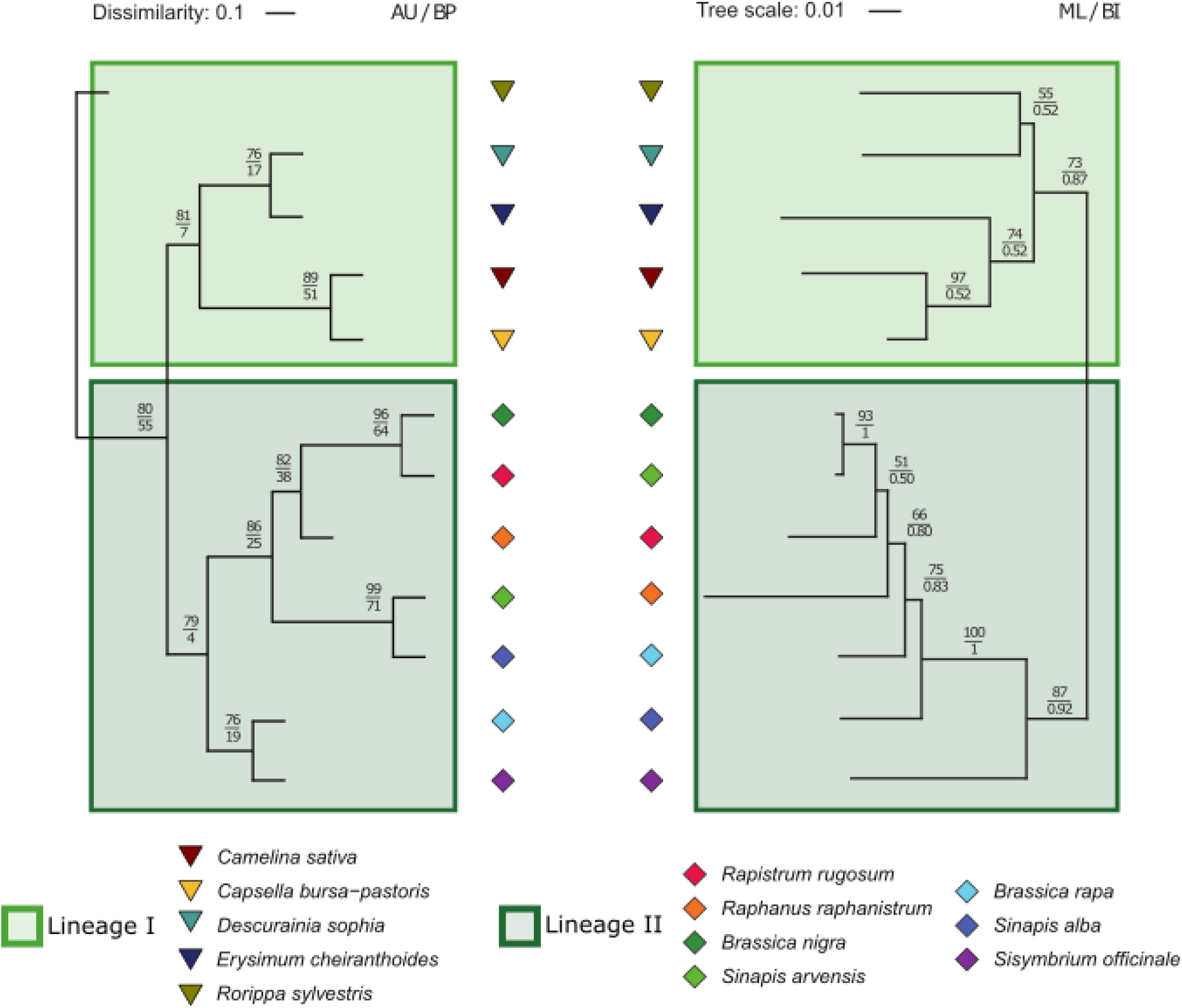
Average multivariate structure of herbivore communities matches with plant phylogeny. Comparison between cluster analysis of the centroids of the log (x + 1) transformed cumulative herbivore abundance data as calculated from PCA coordinates, and the Maximum Likelihood phylogram of Brassicaceae species inferred from ITS sequences. Values of Approximately Unbiased (AU) and Bootstrap Probability (BP) are displayed in the cluster analysis, and Bootstrap support values (BS) > 50% and Bayesian posterior probabilities (PP) > 0.5 are displayed in the phylogram. Scale bars indicate the Euclidean dissimilarity among centroids, and the proportion of sites along each branch, respectively.

### Plant phenotype correlates with herbivore community composition and structure

Herbivore communities are predominantly structured over plant phylogeny through variation in plant-functional traits (Barker et al. 2018). We assessed the variation in functional traits related to plant development across and within plant species and evaluated the role these traits play in structuring herbivore communities. Plant species differed significantly in phenotypic characteristics such as leaf size, number of leaves, plants size and lifetime (Supplementary File 5 Table A), as well as in their multivariate phenotypes (PERMANOVA; df = 11; Pseudo-*F* = 41.5615; *P* = 0.0001). These differences significantly correlated with their phylogenetic dissimilarity (Mantel test: r_m_ = 0.1490; *P* = 0.0010). Phylogenetic signals in specific plant development parameters such as the plant diameter and leaf size were apparent (Supplementary File 5 Table B), whereas the multivariate phenotype was characterized by large variation among all species without consistent differences between species of the two lineages (PERMANOVA; df = 1; Pseudo-*F* = 1.9136; *P* = 0.1449). Pairwise comparisons indicated that the multivariate phenotype of individual plants was significantly different among nearly all plant species (Supplementary File 5 Table C).

Both unconstrained (PCA) and constrained ordination (RDA) indicated that plant lifetime, size of the largest leaf, and plant diameter were often associated with the largest proportion of variation in herbivore communities (Supplementary File 5 Table D – E). The total amount of variation constrained by the plant phenotypic parameters depends on the community subset (Constrained community composition: full: 18.73%; chewing: 18.00%; sap-feeding: 23.03%; Constrained community structure: full: 26.17%; chewing: 20.88%; sap-feeding: 24.91%). The percentage of intraspecific variation in herbivore communities that we could constrain with plant phenotypic parameters was generally low and strongly dependent on the plant species. Interestingly, phenotypic traits better constrained the variation in herbivore communities when taking the abundance of herbivore species into account (Supplementary File 5 Table F – G). These results indicate that the weighted abundance of herbivore species in the community with which plants interact, rather than the identity of the herbivores, correlates more strongly with plant phenotypic parameters.

### Evolutionary models predicting phylogenetic signals

To evaluate which modes of evolution are driving the plant phylogenetic signals in herbivore communities, we tested whether the estimated diversity at the level of plant species and plant individuals was more similar between closely related plants than expected by a model of neutral evolution or a fully random distribution of diversity values (Pagel 1999, Münkemüller et al. 2012). We found that phylogenetic autocorrelation in terms of herbivore richness associated with plant species, estimated independent of branch lengths within the phylogenetic tree, was not significantly different from that expected by neutral or random modes of evolution (Supplementary File 6 Table A). The evolutionary models best fitting the observed distribution of herbivore species richness over plant phylogeny were either fully random models, or models expressing neutral modes of evolution (e.g. Brownian evolution) (Supplementary File 6 Table B). For example, the increased species richness of the sap-feeding herbivore community associated with Brassicaceae species within Lineage II is best explained by a broad model of evolution. In addition, no further phylogenetic signals were found that structure the two Brassicaceae lineages in terms of the richness of communities associated with plant species. Beside the patterns observed for plant species, phylogenetic analysis indicated that plant individuals of closely related species were not more similar in terms of the richness or diversity of herbivore communities than expected by neutral modes of evolution (Supplementary File 6 Table A). This corroborates with the evolutionary models that best fitted the distribution of herbivore species richness associated with plants: i.e. models fitting a neutral mode of evolution or were unable to detect phylogenetic signals (fully random models) (Supplementary File 6 Table B). Whether the observed plant phylogenetic signal in herbivore species diversity across individual plants or plant species emerged as the result of unidirectional evolution and release from herbivore pressure is thus still inconclusive.

## Discussion

By analysing plant phylogenetic signals in insect herbivore communities at the level of individual plants of 12 plant species in two lineages of the Brassicaceae, our study reaches two important conclusions. First, plant phylogenetic signals in the composition and diversity of plant-associated insect communities may be revealed more prominently when focusing on realized interactions from a plant individual perspective rather than describing the full set of interacting species on the plant species level. Second, plant phylogeny predicts the uncertainty of antagonistic interactions at the level of individual plants. This uncertainty is in the proportion of realized interactions per individual plant out of the full potential of interactions with the herbivore pool associated with a plant species. While this proportion is comparable across the two plant lineages, it represents a higher absolute number of interactions for plants in Lineage II. Plants of Brassicaceae Lineage II are attacked by a larger number of herbivore species from a larger species pool, resulting in a higher uncertainty in terms of realized antagonistic interactions compared to plants in Lineage I.

Among the full set of organisms with which a single plant species interacts, phylogenetic relationships have been found to be stronger between plants and their antagonists compared to interactions with their mutualists (Fontaine et al. 2009, Cirtwill et al. 2020). Even though obligate specialists are more readily found in mutualistic interactions, these networks are predominantly composed of more generalized interactions in which pollinators visit flowers of a large number of plants across families (Waser et al. 1996, Thébault and Fontaine 2010). In antagonistic interactions, herbivores are often adapted to specific plant species, genera, or families, resulting in stronger specialization of interaction networks (Fontaine et al. 2009, Thébault and Fontaine 2010). The low level of network specialization for all herbivores interacting with the 12 plant species may be explained by the presence of a unique class of defence chemicals (glucosinolates) in all Brassicaceae (Caballero et al. 2003), which has resulted in dominance of specialist herbivores in Brassicaceae-associated insect communities (Root 1973, Frenzel and Brandl 1998). This causes a large overlap in the composition of the species pools associated with each plant species within the Brassicaceae family (Novotny et al. 2002, Ødegaard et al. 2005, Lind et al. 2015). Only four herbivore species were specific to Lineage II, namely the large cabbage white *Pieris brassicae*, the cabbage whitefly *Aleyrodes proletella*, the marbled yellow pearl *Evergestis extimalis*, and the harlequin bug *Murgantia histrionica*. A single herbivore species, the gall midge *Dasineura sisymbrii*, was exclusively observed on the Lineage I species *Rorripa sylvestris*. These observations may emerge through apparent specialization due to rarity (e.g. *A. proletella*, which was only observed three times in our experiment) or due to context dependency of interactions between plant and herbivore species (Rivera-Hutinel et al. 2012, Chamberlain et al. 2014, Poisot et al. 2015, Costa et al. 2016).

Nevertheless, a plant phylogenetic signal in herbivore communities could be revealed at the lineage level within the Brassicaceae by analysing realized interactions on individual plants. The average diversity, composition, and structure of herbivore communities on individual plants was plant species specific and phylogenetically structured. Thus, plant individuals from closely related species interacted with more similar herbivore communities. Herbivores specializing on plants within the same family are more likely to select plants within a family by similarity in functional traits, which do not necessarily reflect (strong) phylogenetic similarity (Ibanez et al. 2016, Endara et al. 2017). In our study, growth traits partly correlated with the incidence of specific herbivore species on individual plants and acted as a stronger predictor of the weighted abundance of herbivores in realized communities. It is likely that these growth traits covaried with unmeasured but more important (suites of) functional traits such as the plant’s composition of glucosinolates, presence of additional chemical classes such as cardenolides, or production of volatiles, all of which potentially select for some specialization in herbivore interactions (Barker et al. 2018, Blazevic et al. 2020, Züst et al. 2020). Even though plant species may be exposed to a similar pool of potential herbivores, the distribution of these herbivores over individual plants may reveal deeper understanding of phylogenetic signals structuring plant-herbivore communities.

Inferring phylogenetic signals based on herbivore communities on individual plants also revealed a plant-structured phylogenetic signal in uncertainty of attack. Plants in Lineage II seemed to encounter more uncertainty in their interactions compared to plants in Lineage I. This pattern was predominantly driven by differences in both the number of potential and realized interactions on plant individuals. Individuals of Lineage II species were attacked by a more species-rich herbivore community and experienced higher uncertainty in the subset of the full potential pool of herbivores that colonized a plant individual. In addition, plant individuals of the same species had a relatively consistent number of realized interactions but differed in the species composition of this realized herbivore community (i.e. a turnover in herbivore species). The difference in uncertainty of attack may be of major importance for plant defence evolution (Bolnick et al. 2011, Violle et al. 2012, Zytynska and Weisser 2016, Barbour et al. 2018). First, the larger uncertainty of attack could result in stronger selection on induced defences in Lineage II species, including selection for a larger scope of specificity in their responses to cope with the larger pool of potential attackers (Agrawal 2007, Karban 2011). Second, if occurrence of specific herbivore species on individual plants is correlated, the full pool of herbivore species is partitioned in more predictable subsets of realized communities on individual plants. These processes may occur through correlation of herbivore species in response to variation in plant traits, or through priority effects in which presence of one herbivore predicts the course of insect community assembly (Kuppler et al. 2016, Stam et al. 2018). A larger pool of species interacting with a plant species may then result in large variation of subsets of communities on individual plants. These processes may explain the presence and maintenance of genetic variation in defence syndromes in populations, such as the occurrence of distinct plant chemotypes that each harbour a more predictable subset of realized interactions with antagonists (van Leur et al. 2008, Lackner et al. 2019, Patterson et al. 2019). Both hypotheses also requires establishing whether only key herbivore species are selecting on plant defences or whether selection is mediated by direct and indirect interaction in antagonistic communities as a whole (Ohgushi 2005, Poelman and Kessler 2016). In addition, the structuring of herbivore communities over the development of plants may be as important as the compositional structure of herbivore communities in determining the uncertainty of interactions plants experience (Mertens et al. 2020a, Mertens et al. 2020b). However, it remains unknown whether uncertainty in terms of the interactions plants experience is dynamic over the development of plants and how such changes may determine ontogenetic variation in defence strategies. We were not able to identify specific modes of evolution matching the phylogenetic signals we found in the full or average herbivore communities associated with plants. In all cases we found that neutral modes of evolution, or a fully random distribution of diversity measurements over the plant phylogeny best matched our observations. These patterns can be caused by a variety of reasons, including a lack of unidirectional selection on traits affecting interactions with antagonists, differential selection on traits by herbivores and mutualists, an evolutionary arms-race with antagonists, or the under sampling of the phylogenetic tree (Butler and King 2004, Ives et al. 2007, Karinho-Betancourt et al. 2015, Cirtwill et al. 2020). Yet, the plant phylogenetic signal suggests that herbivore community composition on individual plants is derived from plant-herbivore co-evolution.

## Conclusion

We conclude that our findings of a phylogenetic signal in herbivore communities and uncertainty of attack on individual plants are important drivers of evolution of plant defence strategies. We speculate that substantial intraspecific variation in plant-herbivore interactions and the associated uncertainty of attack may maintain genetic variation within plant populations as observed in chemotypes and is likely to select on inducible defence strategies of plants. We thus pledge for a stronger focus on realized interactions between individual plants and herbivore species when inferring phylogenetic signals of interaction partners. This will provide clear opportunities to explore how uncertainty in such realized interactions corresponds with plant defence strategies and will provide novel insights into the drivers of evolution of (plastic) plant defences.

## Materials and methods

### Study system and plant rearing

We monitored the herbivore community associated with 12 annual Brassicaceae species that have overlapping niches and are common in The Netherlands. They belong to two major phylogenetic lineages of the Brassicaceae family (Beilstein et al. 2008) (Supplementary File 1 Table A, Supplementary Fig. 1). Seeds were sown on peat soil (Lentse Potgrond) and germinated under glasshouse conditions. One-week-old sprouts were transplanted into peat soil cubes. One week prior to the start of the field experiment, plants were allowed to acclimatize to field conditions under a roofed shelter. Four-week-old plants were transplanted to the experimental field (mid-May; week 22 of 2016).

### Experimental site and design

The study site was located on the experimental fields of Wageningen University, The Netherlands (51°59′26.5″N 5°39′50.5″E). The experimental fields are embedded in an agricultural and grassland landscape with a variety of brassicaceous herbs, ensuring the presence of a large species pool of insect herbivores part of a typical community on Brassicaceae. We installed a common garden experiment consisting of 120 plots organized in a rectangle of 10 columns by 12 rows. Plant species were randomly assigned to one of the 12 plots within a column, resulting in 10 replicate plots per plant species. Plots measured 3 x 3m and contained nine individual plants in monoculture planted one meter apart. Plots were separated from each other and the field edge by 4-meter-wide grass lanes. To obtain edge uniformity, we planted a strip of *Brassica nigra* (six plants per square meter) around the experimental field. Kites and a meshed fence were placed to prevent damage by vertebrate herbivores. Plots were regularly weeded, and the grass lanes were mown every other week.

### Field observations of herbivores and plant growth

Herbivore communities were monitored on five central plants per plot (i.e. excluding the four corner plants). In cases where a central plant died before the second monitoring round, we monitored one of the corner plants. We recorded naturally occurring herbivores on these plants by weekly counts early in the season and by biweekly counts later in the season. Community development on individual plants was surveyed until seed set. To allow the community to fully develop, insects were identified *in situ* to the highest taxonomic resolution possible (species or family level) (Supplementary File Table B). We aggregated the collected observation data by summing herbivore species abundance per plant over the season, resulting in a singular plant-individual x herbivore-species matrix. This herbivore community matrix could be further divided into a sap-feeding herbivore community matrix and a chewing herbivore community matrix. Previous studies have shown that feeding guilds differ in their interaction with Brassicaceae, are different in their level of host specialization, and are governed by different processes at both regional and local scales (Lewinsohn et al. 2005, Soler et al. 2012). Moreover, parameters describing diversity of the herbivore community may be strongly affected by the biology of the herbivore species involved. For example, the evenness of herbivore communities is likely to be biased to emphasize the role of aphids in herbivore communities, obscuring variation across plants or plant species in terms of their interactions with chewing herbivores, which arguably also impose a substantial fitness cost on plants. Determining whether the uncertainty in the interactions plants experience is mainly driven by variation in diversity of one of the two respective feeding guilds or by variation in diversity of the herbivore community in its entirety may reveal which underlying mechanisms or plant strategies may play a more prominent role in shaping the uncertainty of interactions.

In addition to arthropod observations, we recorded a set of phenotypic parameters for all visited plants during each monitoring round: plant height (measured from the ground to the top of the plant), diameter (measured as the distance between the two most distal leaves), length of the largest leaf, number of true leaves, and number of flowering branches. The plant traits that were measured can readily be hypothesised to affect the herbivore community, as they determine the apparency of plants (e.g. plant height) or the availability of specialised niches (e.g. number of flowering branches). Importantly, these measurements are non-destructive and ensure a minimal disturbance of the assembly of the plant-associated herbivore community. In the subsequent analyses, we used the maximum observed parameter values for each plant individual to represent its specific phenotype.

### Reconstruction of plant phylogeny

ITS sequences of the 12 brassicaceous species and the outgroup *Aethionema arabicum*, a sister species of the core Brassicaceae, were retrieved from the BrassiBase website (https://brassibase.cos.uni-heidelberg.de). Multisequence alignments were obtained by MAFFT v7 using the iterative refinement method FFT-NS-i with a gap opening penalty of 1.0. Unreliable alignment regions were detected with GUIDANCE2 using 100 bootstraps at default thresholds. Residues and columns with a confidence score > 0.650 were removed. Phylogenetic relationships were inferred by maximum likelihood (ML) and Bayesian methods. ML analyses were computed with W-IQ-TREE using the best-fit model (SYM+G4) selected by ModelFinder and 1000 bootstrap replicates. Bayesian interference analysis was conducted with MrBayes 3.2.6 using the GTR substitution model under default priors. Chains were run for 100,000 generations and trees were sampled every 100 generations. The initial 1000 trees were discarded as burn-in, and posterior probabilities (PP) were calculated from the remaining replicates. Phylogenetic trees were drawn and edited in iTOL 4.4 (Supplementary Fig. 1).

### Data analysis

To test for plant phylogenetic signals in herbivore communities on the level of plant species or plant individuals, we explored the incidence-based as well as the abundance-based community dataset and its subsets (focusing on the community of sap-feeding herbivores or chewing herbivores). We summed all herbivore observations per plant over the season into a singular plant individual x herbivore species matrix. From this matrix we derived matrices that present observations at the biological level of plant species, or which are transformed to adjust for potential biases that are inherent to the biology of the herbivore species in our community. An overview of the different matrices and the analyses in which they were used is presented in the supplementary information (Table 1).

Framing diversity in the context of interactions among species can provide insight in the structure and niche dimensions of plant-associated herbivore communities. To this end, we calculated a set of interaction network metrics: Network connectivity C (Delmas et al. 2019), nestedness ADDIN EN.CITE (weighted NODF; Almeida-Neto et al. 2008, Almeida-Neto and Ulrich 2011) and specialization (H2’ and d_i_’) of the plant species-herbivore interaction networks (Table 1). We tested the network metrics by comparing them to two separate null models based on the Patefield (Patefield 1981) and on the Vaznull algorithms (Vazquez et al. 2007), respectively. We generated 999 random networks for each of the algorithms and used one-sample t-tests with our observed network descriptor as reference to estimate significance (Blüthgen et al. 2008, Flores et al. 2011).

We then calculated three widely used community diversity indices, namely species richness, Shannon diversity and Simpson diversity for each plant species and on the level of each plant individual (Heip et al. 1998) (Table 1). In addition, we compared the proportion of realized interactions by relating the richness of the species pool that could interact with a plant species to the average richness of herbivores observed on individual plants of that species (Whittaker’s β diversity). This metric expresses the number of times the richness of the species-associated herbivore pool is greater than the herbivore richness observed for plant individuals of this species. These community properties were tested for phylogenetic signals by applying (generalized) linear (mixed) effect models ((G)L(M)M). To verify the completeness of our survey and assess whether uneven sampling efforts would affect our conclusions, we evaluated sampling completeness (Chao and Jost 2012), explored sample-based species rarefaction curves and compared diversity measures at interpolated and, where relevant, at extrapolated estimations of sample completeness (Supplementary Figure 6, Supplementary File 3 Table A) (Gotelli and Colwell 2001). Herbivore communities were further characterized by the dissimilarity in herbivore species composition between plant individuals of the same species (expressed by multivariate Sørensen β diversity). Analysing multivariate β diversity is complementary to the analysis of species richness, as this index emphasizes differences in species identities among the plant-associated herbivore communities (Anderson et al. 2011). The Sørensen dissimilarity has a direct correspondence with multivariate dispersion in community composition (Anderson et al. 2006) and can be decomposed into a turnover component (i.e. dissimilarity due to species replacement), and a nestedness component (i.e. dissimilarity due to loss or gain in species richness) (Baselga 2010, 2012).

We used multivariate ordinations (non-metric multidimensional scaling) of the herbivore community on individual plants to explore the variation in composition and structure of herbivore communities associated with plants across the different plant species and lineages. We assessed community composition based on the Sørensen dissimilarity matrix (NMDS: three ordinal dimensions, stress = 0.18) and community structure by the Bray-Curtis dissimilarity matrix (NMDS: three ordinal dimensions, stress = 0.19) (Table 1). We then estimated the overall dissimilarity in herbivore communities explained by plant species and lineages by permutational analyses (PERMANOVA) (Anderson 2001). To ensure valid permutation of communities, we specified the nested structure of our experiment in the permutational design. Statistical significance was assessed via 999 random permutations. Post-hoc comparisons between plant species were made by running a separate PERMANOVA analysis for each comparison and limiting the proportion of type I errors by false discovery rate control (FDR) (Verhoeven et al. 2005).

We then applied mantel tests to quantify the correlation between the phylogenetic similarity of plant species and the similarity in herbivore communities on plants (Mantel 1967, Legendre and Legendre 1998). To visually compare the relation between the similarity of herbivore communities and relatedness of plant species, we ran an unconstrained principal component analysis (PCA) on the abundance-based community data (Table 1, Matrix C) and calculated the multivariate coordinates of the centroids of each of the plant species. We then used these coordinates in a cluster analysis (complete linkage, euclidean distance), and verified the clusters by using a multiscale bootstrap procedure (100 bootstraps) (Suzuki and Shimodaira 2006).

The herbivore communities were tested for plant-phylogenetic signals by applying Mantel tests, evaluating the correlation between the dissimilarity among plants in terms of their ITS sequences and the dissimilarity among plants in terms of their herbivore communities. We then estimated the contribution of differences in herbivore richness and turnover of herbivores to variation in herbivore communities within plant species (expressed by multivariate β diversity and its components). We then related variation in herbivore communities and its subsets to the phenotypic traits measured for plant individuals. To allow comparison of different traits, we scaled each of the traits to zero mean and unit variance. As similarity in plant phenotypes are likely to be higher for closely related species (Blomberg and Garland 2002), we first tested the correlation between phenotypic traits and phylogenetic relatedness by applying a mantel test. We then ran an unconstrained PCA ordination on our community data, followed by post-hoc regression of the three most important PCA axes by the plant-phenotypic traits (Legendre and Legendre 1998) (Table 1). In an alternative approach, we constrained the variation in the herbivore community with the measured plant phenotype by applying a stepwise redundancy analysis (RDA) procedure with 999 random permutations per step (Van den wollenberg 1977, Šmilauer and Lepš 2014). To determine whether the relation between plant phenotype and the associated communities differed among plant species, we repeated the stepwise RDA procedure for each plant species separately.

Finally, we evaluated which evolutionary models best predict the phylogenetic signals in herbivore community diversity and uncertainty of attack in Brassicaceae by calculating the phylogenetic autocorrelation represented by Abouheif’s C_mean_ (Münkemüller et al. 2012, Keck et al. 2016). To complement this analysis, we fitted a set of models simulating different modes of evolution, taking the branch length of the phylogeny into account These included a model expressing Brownian evolution, a model estimating phylogenetic signal by Pagels Lambda (Pagel 1999), and a fully random model based on white noise (Münkemüller et al. 2012). By comparing AICc values, we then selected the evolutionary model that fitted the distribution of estimated diversity values over the plant phylogeny best. As branch lengths can greatly affect the fit of these evolutionary models, we also analysed the fit of the evolutionary models separately for the two lineages.

All statistical analyses were performed using R (v3.2.4) (R Core Team 2014) packages nlme (Pinheiro et al. 2012), lme4 (Bates et al. 2015), emmeans (Lenth et al. 2019), vegan (Oksanen et al. 2012), BiodiversityR (Kindt and Coe 2005), bipartite (Dormann et al. 2008, Dormann et al. 2009, Dormann 2011), ape (Paradis et al. 2004), Geiger (Harmon et al. 2008) and pvclust (Suzuki and Shimodaira 2006).

## Data availability

The data reported in this paper has been deposited at Dryad (https://doi:10.5061/dryad.4j0zpc88c).

## Acknowledgments

We acknowledge Freek Bakker for advice on phylogenetic analysis, Marcel Dicke for feedback on our manuscript, and the staff of Unifarm for setting up and maintaining the experimental fields. This project was supported by the European Research Council (ERC)

Under the European Union’s Horizon 2020 research and innovation program (Grant agreement No. 677139 to E.H.P.).

## Author Contributions

EHP conceived the study and designed the experiment, DM collected the data and performed the statistical analyses, and KB constructed the phylogenetic tree. All authors contributed substantially to the writing of the manuscript.

## Additional Information

**Supplementary Information** is available for this paper

---

**Supplementary Fig. 1.**
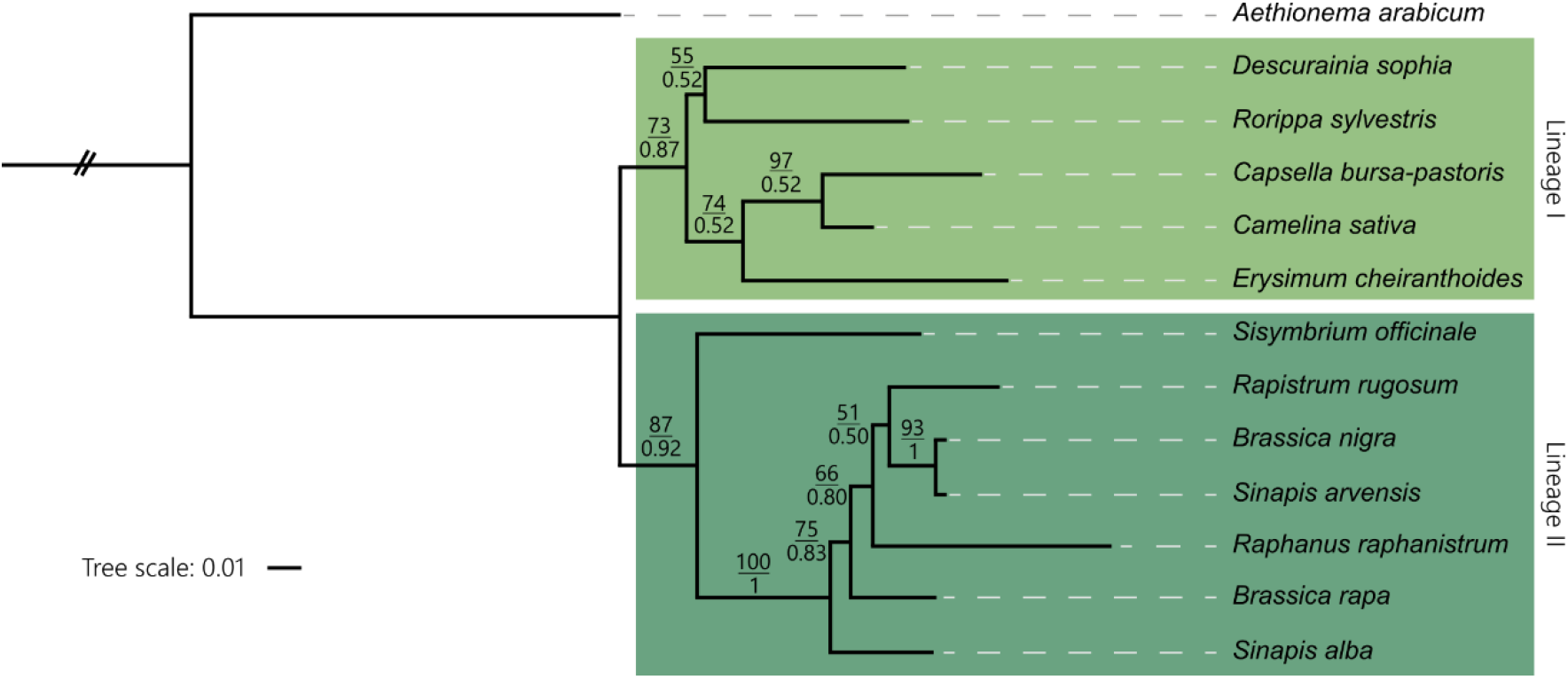
Maximum Likelihood phylogram of brassicaceous species inferred from ITS sequences. The phylogenetic tree was rooted to the sister species. Bootstrap support values (BS) > 50% and Bayesian posterior probabilities (PP) > 0.5 are displayed at branches. Scale bar indicates the number of nucleotide substitutions per site.

**Supplementary Fig. 2.**
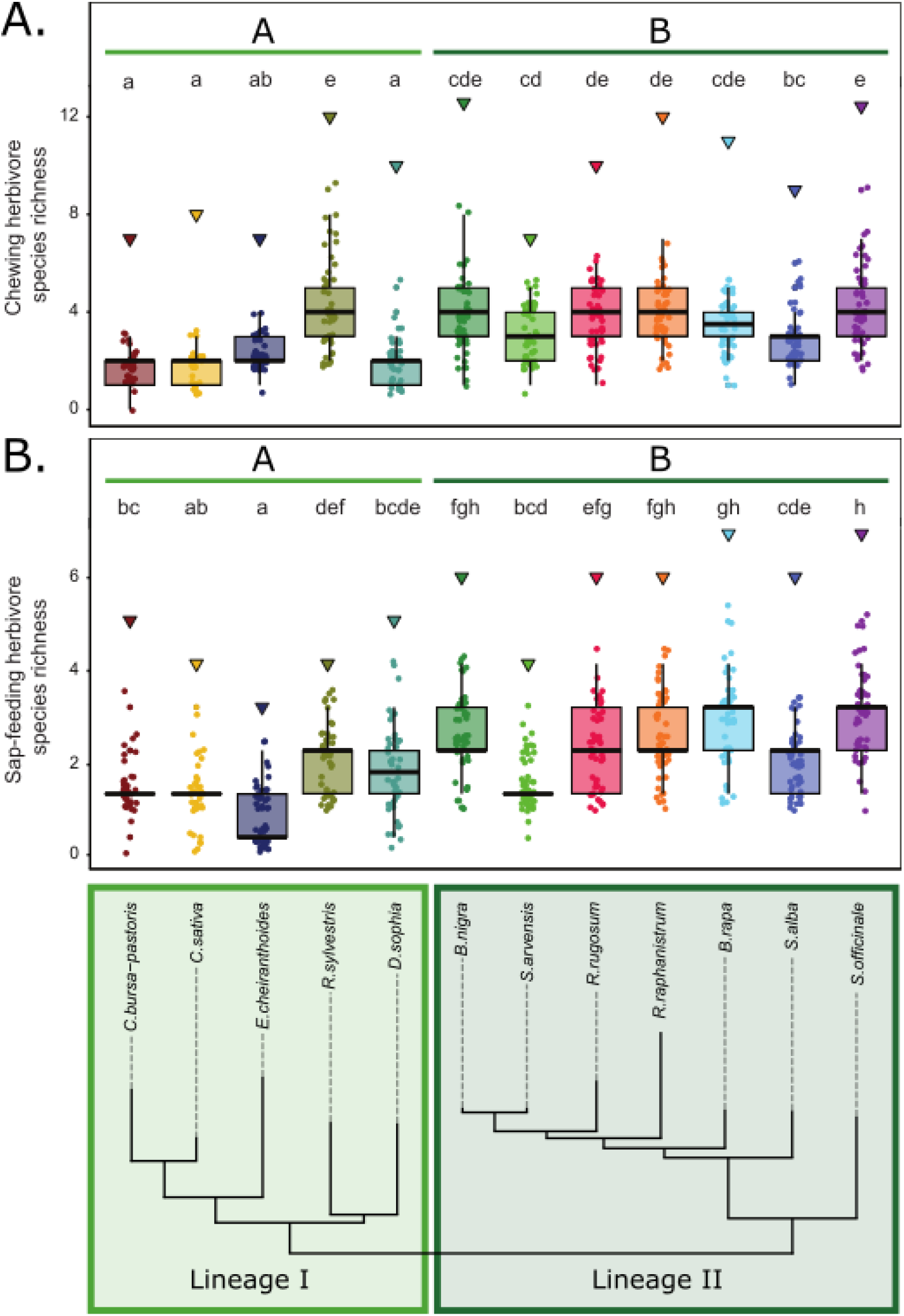
Herbivore species richness is dependent on plant species and structured by plant phylogeny. (A) Species richness of the chewing herbivore community and (B) species richness of the sap-feeding herbivore community observed across plant phylogenetic lineages (lower panel). Triangles depict the total number of herbivore species of the respective feeding guild, associated with the total number of individuals per plant species. Dots represent the number of herbivore species observed on plant individuals. Box – whisker plots summarize the variation in observed richness. Statistical analyses were performed by applying Linear Mixed Models (LMM) with species or phylogenetic lineage as explanatory factors and including plot and, when estimating the diversity for phylogenetic lineages, plant species as random factors in our models. To account for heterogeneity of variance, we allowed the variance to be different for the different species or lineages in our model. Different letters indicate significant different means (*P* < 0.05), adjusted for multiple testing by Tukey HSD. Significant differences across lineages (plant species grouped by the coloured horizontal bars) are indicated with capital letters. Statistical analyses were performed separately for the different panels

**Supplementary Fig. 3.**
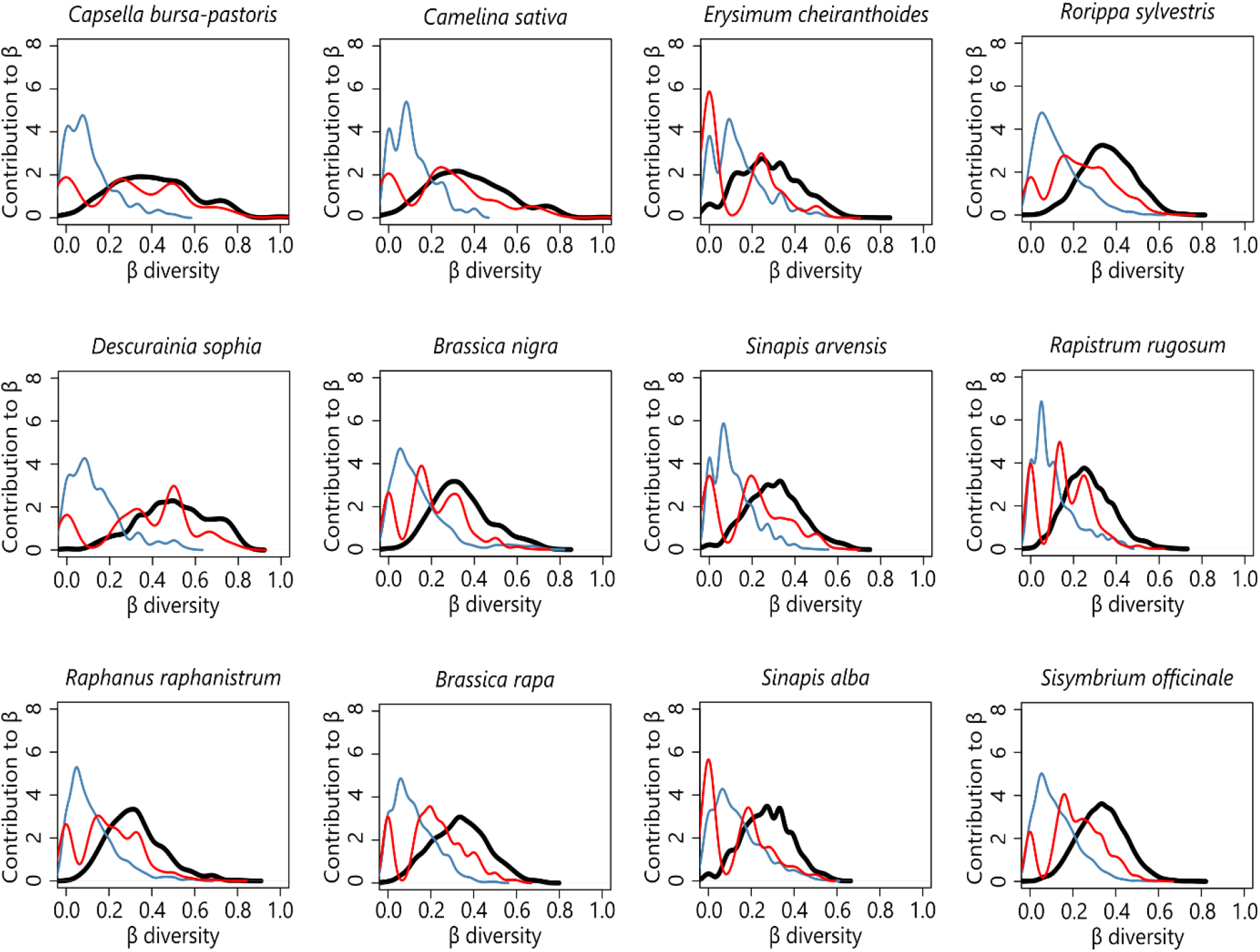
Intra-specific variation in species composition is similar across plant species and predominantly caused by differences in herbivore species identity. Intra-specific variation is represented by the Sørensen dissimilarity (i.e. multivariate β diversity; black line). β diversity can be partitioned in variation caused by differences in species identity (red line) and variation caused by differences in the number of species between plant individuals (blue line).

**Supplementary Fig. 4.**
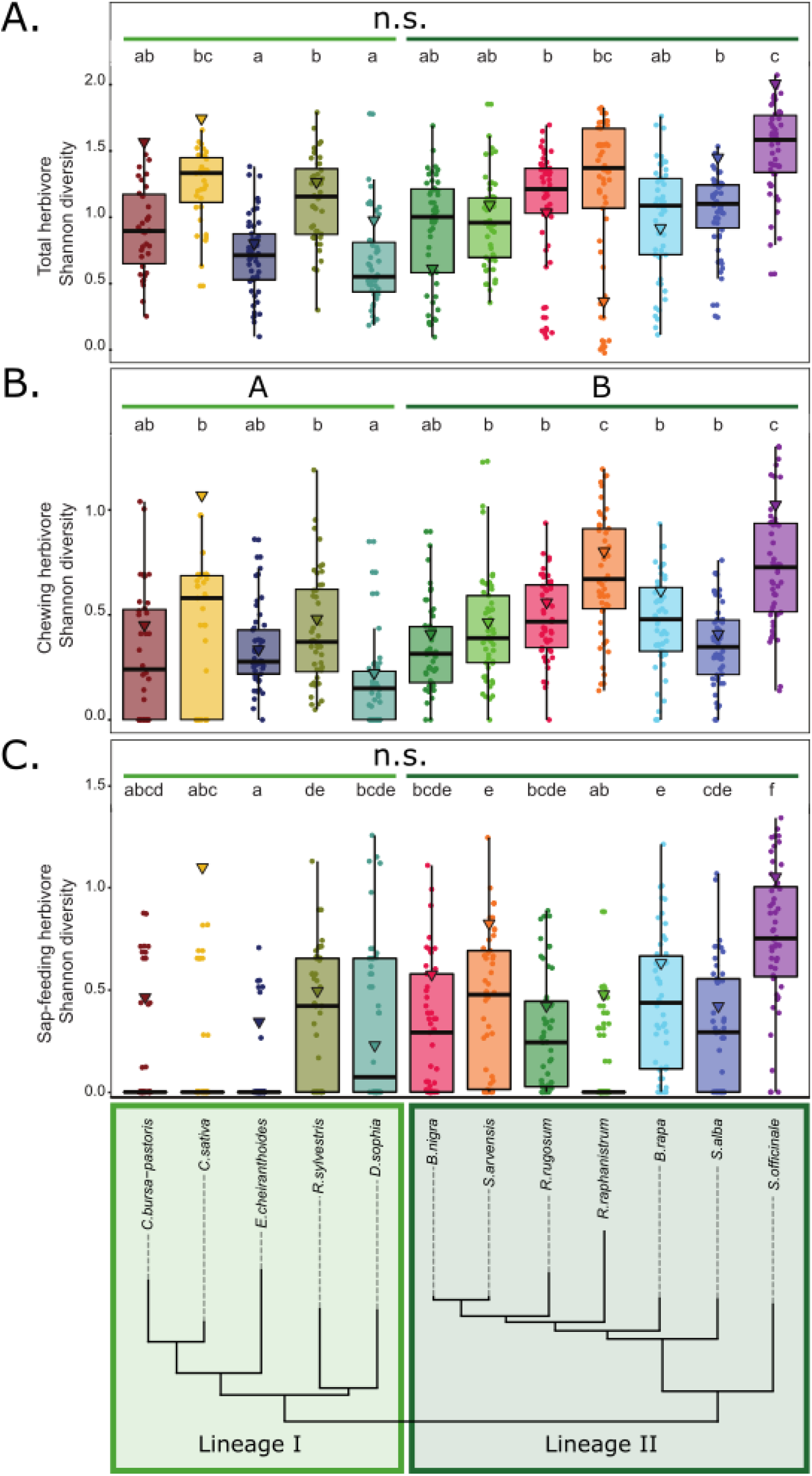
Shannon diversity of herbivore communities is dependent on plant species and structured by plant phylogeny. (A) Shannon diversity of the full herbivore community (B) Shannon diversity of the chewing herbivore community, and (C) Shannon diversity of the sap-feeding herbivore community associated with plant species from two phylogenetic lineages in the Brassicaceae (lower panel). Triangles depict the Shannon diversity of the total herbivore community or the diversity of the respective feeding guild, associated with the total number of individuals per plant species. Dots represent the observed Shannon diversity on plant individuals. Box – whisker plots summarize the variation in Shannon diversity. Statistical analyses were performed by applying Linear Mixed Models (LMM) with species or phylogenetic lineage as explanatory factors and including plot and, when estimating the diversity for phylogenetic lineages, plant species as random factors in our models. To account for heterogeneity of variance, we allowed the variance to be different for the different species or lineages in our model. Different letters indicate significant different means (*P* < 0.05), adjusted for multiple testing by Tukey HSD. Significant differences across lineages (plant species grouped by the coloured horizontal bars) are indicated with capital letters. Statistical analyses were performed separately for the different panels.

**Supplementary Fig. 5.**
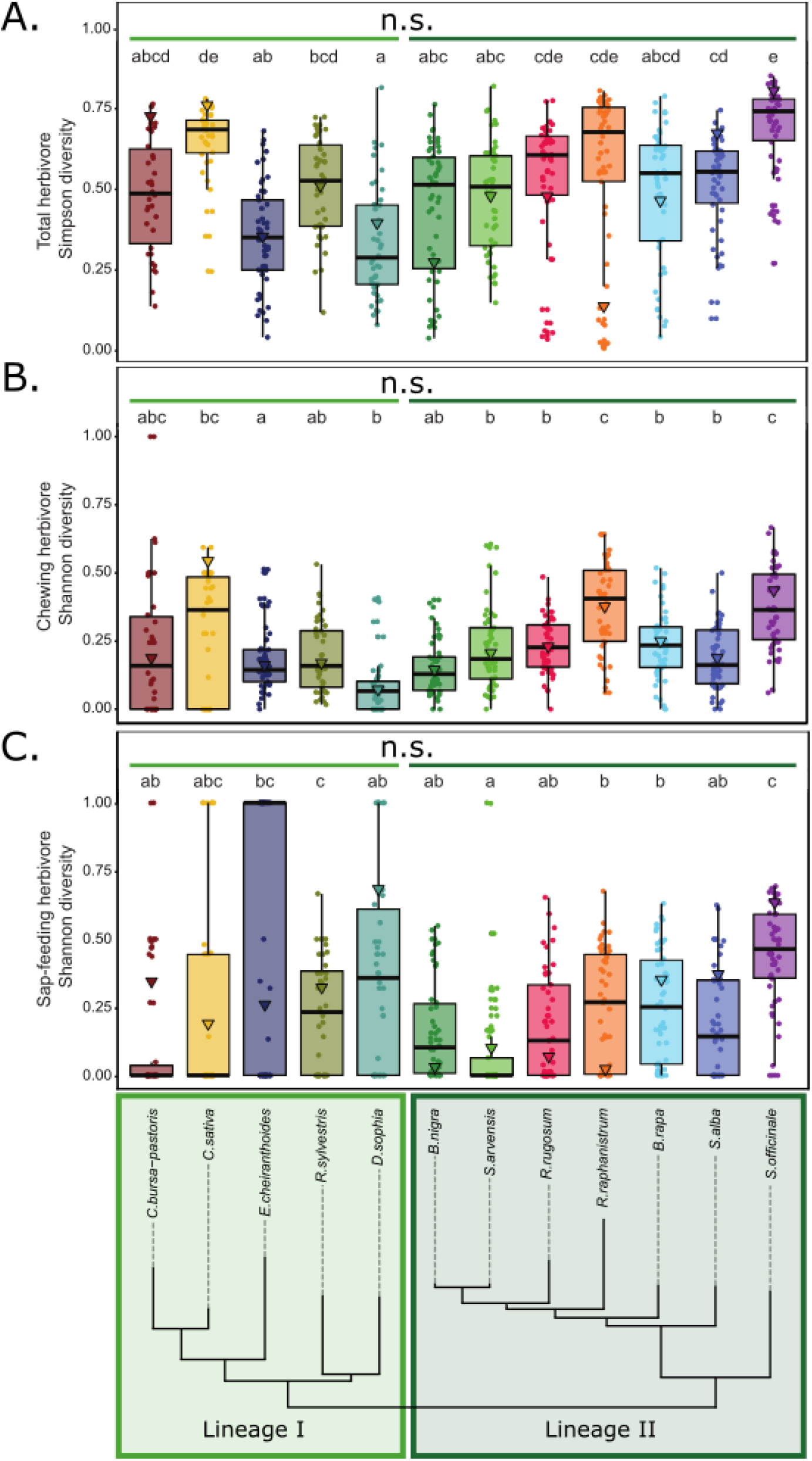
Simpson diversity of herbivore communities is dependent on plant species and structured by plant phylogeny. (A) Simpson diversity of the full herbivore community (B) Simpson diversity of the chewing herbivore community, and (C) Simpson diversity of the sap-feeding herbivore community associated with plant species from two phylogenetic lineages in the Brassicaceae (lower panel). Triangles depict the Simpson diversity of the total herbivore community or the diversity of the respective feeding guild, associated with the total number of individuals per plant species. Dots represent the observed Simpson diversity on plant individuals. Box – whisker plots summarize the variation in Simpson diversity. Statistical analyses were performed by applying Linear Mixed Models (LMM) with species or phylogenetic lineage as explanatory factors and including plot and, when estimating the diversity for phylogenetic lineages, plant species as random factors in our models. To account for heterogeneity of variance, we allowed the variance to be different for the different species or lineages in our model. Different letters indicate significant different means (*P* < 0.05), adjusted for multiple testing by Tukey HSD. Significant differences across lineages (plant species grouped by the coloured horizontal bars) are indicated with capital letters. Statistical analyses were performed separately for the different panels.

**Supplementary Fig. 6.**
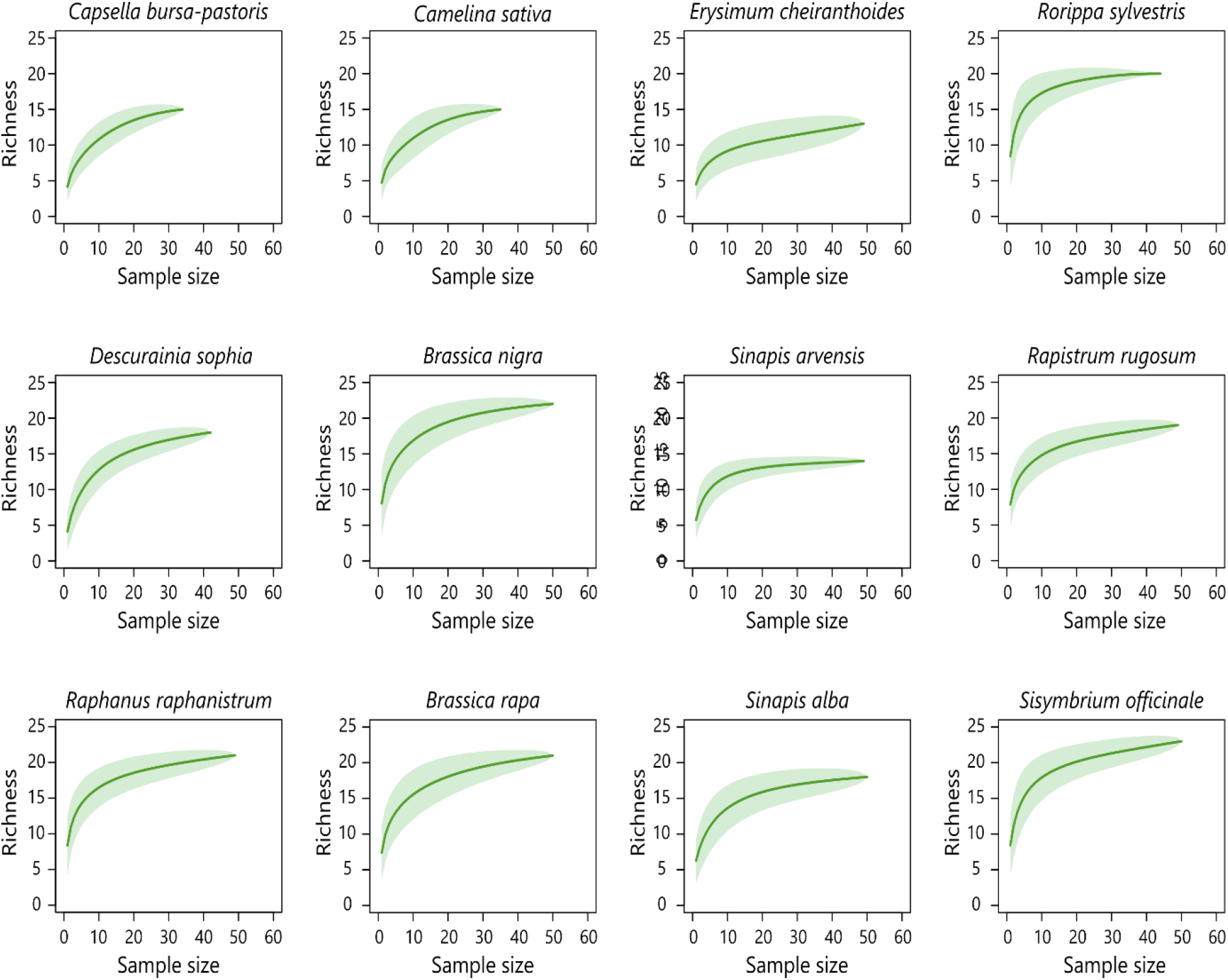
Sample-based species rarefaction curves indicate high sample completeness (79.64% or higher). Curves representing mean species richness (solid line) and variation of repeated re-sampling (shaded area). Curves reaching an asymptote are indicative of high levels of sampling completeness.

